# Do El Niño events beget generations of reproductively deficient adults?

**DOI:** 10.1101/594226

**Authors:** Santiago Ortega, Cristina Rodríguez, Hugh Drummond

**Affiliations:** Posgrado en Ciencias Biológicas, Universidad Nacional Autónoma de México; Departamento de Ecología Evolutiva, Instituto de Ecología, Universidad Nacional Autónoma de México, 04510, Ciudad de México, México

**Keywords:** Adverse environments, cohort effects, resilience, lifetime reproductive success, El Niño Southern Oscillation, viability selection, self-selection

## Abstract

1. Warm ocean waters during El Niño events deplete primary productivity, with cascading effects through the food chain that profoundly affect many marine and terrestrial species, commonly increasing adult mortality and offspring starvation. With global warming, events will double and increasingly threaten the depletion or extinction of some animal populations.
2. Because adverse environments experienced during infancy generally induce reproductive deficits in adulthood, El Niño events are also expected to affect animals born during them, engendering generations of adults with reduced reproductive potential and exacerbating demographic impacts.
3. We made the first test of this idea, using the blue-footed booby, a piscivorous apex predator of the eastern tropical Pacific Ocean. Surprisingly, detailed monitoring of 18 generations over a 27-year period detected no deficits in the annual breeding success, offspring viability, lifespan or lifetime reproductive success of generations of adults born during El Niño years.
4. These results testify to remarkable developmental resilience extending across the lifespan. However, there was evidence that this resilience was supported by two mechanisms of quality control of adult generations from El Niño years.
5. First, viability selection on nestlings and fledglings was more severe for El Niño birth cohorts than ordinary cohorts.
6. Second, in El Niño years, adult boobies self-selected for breeding. There was no increase in the proportional representation of either high quality breeders or breeders in their peak years (middle-age), but there was an increase in old-young adult pairings, which in this population produce the most viable fledglings.
7. The blue-footed booby appears to be immune to the expected developmental impact of El Niño on the reproductive value of adult generations. The susceptibilities and resilience of other species need to be explored, to better predict the demographic impacts of this accelerating climatic oscillation.

## Introduction

With global warming, our planet’s most dramatic short-term climate oscillation, El Niño Southern Oscillation (ENSO), is predicted to increase in intensity, with extreme events doubling in frequency in the near future (Cai et al., 2014; Ham, 2018; Wang et al., 2017). This raises urgent questions about the ability of animal populations of the eastern Pacific Ocean to withstand its accelerating demographic impacts. During El Niño events, separated by 2-7 year intervals (Zhang & Zhao, 2015), warm ocean waters deplete primary productivity, with broadly deleterious consequences throughout the food chain. Impacts on marine vertebrates in the reproductive season, especially seabirds, include physiological suppression and delay (Guerra, Fitzpatrick, Aguilar, & Venables, 1988), reduced adult survival and reproduction, nest abandonment, reduced offspring growth and survival, population decline and changes in distribution (Hays, 1986; Hodder & Graybill, 1985; Le Boeuf & Reiter, 1991; Tovar, Guillen, & Cabrera, 1987; Valle, Cruz, Cruz, Merlen, & Coulter, 1987); with increased frequency of warm water events limiting population recovery and increasing risk of extinction (Vargas, Harrison, Rea, & Macdonald, 2006; Vargas et al., 2007).

Another likely consequence, with demographic implications, is diminished reproductive value of adults born in El Niño years. No study has addressed the possibility of El Niño cohort effects, but numerous experiments have shown that early-life environmental conditions usually have developmental effects on physiological, behavioural, morphological and life history traits of vertebrates during adulthood (Grace, Martin-Gousset, & Angelier, 2017; Haywood & Perrins, 1992; Mumme, Bowman, Pruett, & Fitzpatrick, 2015; Nettle et al., 2017; Plard et al., 2015). Furthermore, in mammals, natural early-life adversity has been linked to harmful consequences on fitness components and senescence rates (e.g. Nussey, Kruuk, Morris, & Clutton-brock, 2007; Pigeon, Festa-Bianchet, & Pelletier, 2017; Plard et al., 2015), and in 5 of 9 bird species studied in nature, adults that experienced poor weather, habitats, prey availability, parental care or late fledging in the natal year showed reduced lifetime reproductive success (review in Drummond & Ancona, 2015). There is also experimental and descriptive evidence for intergenerational effects: negative impacts of nutritional and social stress in infancy on the quality and viability of the infant’s eventual offspring (Burton & Metcalfe, 2014; Drummond & Rodríguez, 2013; Naguib & Gil, 2005).

These findings raise the expectation that in many species, generations of adults from El Niño birth cohorts will show survival or fitness deficits or produce offspring of low quality, at some stage of their lifetime. However, in wild animals such effects are sometimes mitigated in one or both sexes by simple developmental resilience or by making life-history adjustments that seemingly neutralize fitness penalties (e.g. Ancona & Drummond, 2013; Cartwright, Nicoll, Jones, Tatayah, & Norris, 2014; Dmitriew & Rowe, 2007; Douhard et al., 2016). Thus, in some species, adults exposed to warm ocean waters in their natal year might sidestep impacts on their lifetime reproductive success, although complete evasion of fitness impacts of early-life environmental stress has not been documented for any long-lived animal.

Our case study of a population of blue-footed boobies (*Sula nebouxii*) tested for effects of ENSO conditions (mean water temperature in the 8-month breeding season) in the natal year on the quality of adult generations by comparing the lifetime reproductive performance of recruits from 18 birth cohorts. This long-lived (up to 25 years), socially monogamous apex predator of the eastern tropical Pacific Ocean lays clutches of 1-3 eggs, but often raises fewer chicks when underfeeding triggers siblicidal brood reduction. When an El Niño event depletes populations of its fish prey, circulating corticosterone increases in females (Wingfield, Ramos-Fernandez, Nuñez-de la Mora, & Drummond, 1999), mortality of young adults increases, fewer boobies breed, breeding is delayed, clutch and brood sizes are smaller, hatching fledging and breeding success decline, and nestlings grow more slowly (Ancona, Sánchez-Colón, Rodríguez, & Drummond, 2011; Kiere & Drummond, 2016; Oro, Torres, Rodríguez, & Drummond, 2010). Nonetheless, previous analysis of the first ten years of life suggested that male and female recruits from El Niño birth cohorts largely escape developmental impacts: they recruited at an earlier age, bred less frequently (skipped more years) and matched the survivorship and accumulated breeding success (production of fledglings) of recruits born under favourable cold water conditions (Ancona & Drummond, 2013). However, these observations do not demonstrate full equivalence to recruits from cold-water cohorts because fitness penalties could manifest after the first 10 years or in other fitness components such as rate of reproductive senescence, adult lifespan or offspring quality. To determine whether El Niño events impact the reproductive value of adult generations, fitness accounting across the whole lifespan of recruits is required. This was possible for the focal population because, with rare exceptions, fledglings nest for the first time (mostly at ages 2-5 years) close to where they hatched, then nest close to their first nest site during the rest of their lives (Kim, Torres, Domínguez, & Drummond, 2007).

We also tested for two largely unstudied filtering processes which theoretically could buttress populations against demographic impacts of adverse natal environments and mask their developmental impacts. The first is self-selection for breeding in adverse conditions by high quality breeders able to confer their own quality on offspring. In support of this idea, there is some evidence that in stressful circumstances animals that are in poor condition, very young or very old may skip breeding, trading reproduction for survival in the service of lifetime reproductive success (Cubaynes, Doherty, Schreiber, Gimenez, & Jr, 2010; Goutte, Kriloff, Weimerskirch, & Chastel, 2011; Skjaeraasen et al., 2012), and resulting in an increase in the proportion of high quality individuals among breeders. The second is increased viability selection in adverse conditions: differential survival and recruitment into the breeding population of high quality offspring (cf. Garratt et al., 2015; Mojica & Kelly, 2010).

## Materials and methods

### Study population and data collection

Demographic data were collected between 1989 and 2018 during lifelong annual monitoring of individual blue-footed boobies (*Sula nebouxii)* on Isla Isabel, Nayarit (21°53’N, 105°54’W), off the Pacific coast of Mexico (Drummond, Torres, & Krishnan, 2003; Kim et al., 2007).

Breeders were individually identified by bands fitted at fledging. Throughout each reproductive season (roughly February through July) all pairs in two study areas were monitored by recording their nest contents every 3-6 days through fledging (details in Drummond et al., 2003). Nests in the study area and within 20 m of its borders, which correspond with natural borders of the colony, are too conspicuous on the beach and forest floor to escape detection by the team of two monitors (Drummond & Rodríguez, 2015). This monitoring protocol allowed detection of nearly all recruitment and breeding of the focal fledglings through 2018.

For all analyses of breeders, samples included all members of the 18 birth cohorts (1989-2009) that recruited by age 6 years (the age by which 87% of fledglings recruit), thereby excluding individuals whose early nesting may have been missed. We analysed annual breeding success (number of 70-d old fledglings produced, regardless of brood size), fledgling viability (proportion of fledglings that recruited), adult lifespan (age at the last observed breeding), lifetime reproductive success (total recruits produced), fledging success (proportion of nestlings that managed to fledge), and recruitment (whether an individual was resighted as a breeder; 1-0 variable), separately for females and males because pair mates are not statistically independent. Because nestlings and fledglings are not sexed until they return to the colony as breeders, the analyses of fledging success and recruitment were not separated by sex.

The sample for analysis of annual breeding success included all breeders from the 18 cohorts (2143 male and 1893 female boobies); the sample for analysis of fledgling viability included the offspring produced by all breeders from those cohorts that fledged at least one chick at any age (1646 male and 1329 female breeders), and excluded fledglings born after 2013 because their recruitment could not be adequately scored (they were monitored for less than 6 years). The sample for analyses of adult lifespan and lifetime reproductive success included only the 1319 male and 1105 female breeders that died before 2013, allowing each of their fledglings at least 6 years to recruit during annual monitoring. Death was assumed when a breeder was absent during three consecutive breeding seasons; in the cohorts of 1989-1995, only 5-6% of breeders were ever resighted after three consecutive absences. For males and females, respectively, ages of first reproduction (mean ± SD) were 4.14 ± 0.95 and 3.54 ± 0.98 years, and adult lifespans were 8.05 ± 3.49 and 8.02 ± 3.99 years (range 1 to 22 years for both sexes).

Self-selection for breeding in warm water conditions was tested in the 4036 males and females of the 18 cohorts that recruited by looking for increase in the proportional representation of individuals of above average quality, favourable ages (10-14 years) or from cool water (La Niña) natal years (lest this confer any long-term benefit). Individual quality was expressed as the mean of all z-normal standardized annual breeding success values (fledglings produced) over each individual’s lifespan.

Increased viability selection on warm water cohorts was tested by analysing the proportion of nestlings that managed to fledge each year and the recruitment probability of each year’s fledglings. For fledging success, we used data from one-, two-, and three-chick broods (sample sizes: 7630, 6975, and 1326, respectively) reared between 1989 and 2018; for recruitment, we included all 10604 fledglings produced between 1989 and 2013 for which body measures were available and recruitment could be adequately scored. The ratio between weight (grams) and ulna length (millimetres) at age 70 days was used as an indicator of body condition at fledging. This ratio accounts for the sexual size dimorphism of the boobies; at 79 days old, females weigh approximately 26% more than males, and their ulnas are 10% longer (Torres & Drummond, 1999). To remove year effects, individual body condition was calculated as the z-normal standardized annual body condition at fledging of the colony. Boobies weighed (mean ± SD) 1593.51 ± 256.2 grams at fledging with an ulna length of 205.57 ± 12.98 millimetres.

ENSO conditions during the reproductive season in which a focal breeder was born were expressed as the average of the 8 monthly (December to July) sea surface temperature anomalies (SSTA) at the closest oceanographic station, 55 km from the colony, obtained from the IRI Data Library http://iridl.ldeo.columbia.edu/SOURCES/.NOAA/.NCEP/.EMC/.CMB/.GLOBAL/.Reyn_SmithOIv2/.monthly/.ssta/. Niño and La Niña years were characterized by mean SSTA of at least +0.5°C or −0.5°C, respectively, using the criteria of NOAA, obtained from the El Niño and La Niña Alert System https://www.climate.gov/news-features/understanding-climate/el-ni%C3%B1o-and-la-ni%C3%B1a-alert-system. The Southern Oscillation Index (SOI) was not used because SSTAs are more closely related to the breeding parameters and age at first reproduction of this population (Ancona & Drummond, 2013; Ancona et al., 2011).

### Statistical analysis

All analyses were performed in R statistical environment (R Development Core Team, 2018). All variables were standardized prior to model fitting to minimize collinearity of linear and quadratic terms and to facilitate the interpretation of the relative strength of parameter estimates (Cade, 2015; Grueber, Nakagawa, Laws, & Jamieson, 2011). Because high or even moderate collinearity can inflate standard errors of coefficient estimates and make genuine effects harder to detect (Zuur, Ieno, & Elphick, 2010), the variance inflation factor (VIF) was calculated for each variable in every hypothesis-based model, so that variables with VIF values higher than 2 could be excluded from the final analyses (none were). All VIF values are reported in the supplementary materials (Supporting information). Models were fitted with a Gaussian error distribution and an identity link function. Variable standardization was carried out using the *rescale* function in the R package *arm* (Gelman et al., 2016). We used the *lmer* function in the package *lme4* (Bates, Maechler Martin, & Walker, 2016) to build the LMMs; LMs were built using the built-in function *lm* in R. Model selection was carried out with *model.sel* function in the MuMIn package (Bartón, 2016)35, *mod.avg* function was used to average models within a 95% confidence interval (CI; Burnham, Anderson, & Huyvaert, 2011), and *confint* function to calculate the CI of each parameter.

To assess the effects of recruits’ early-life conditions on their annual breeding success and their fledglings’ viability, we first constructed a base model (a LMM accounting for previously demonstrated effects in the species; *sensu* Panagakis, Hamel, & Côté, 2017) for each analysis; this contained the linear and quadratic expression of breeder age and laying date, with breeder identity as random effect. For annual breeding success, SSTA in each reproductive year was also included in base models. The linear expression of year of birth was included in every model, irrespective of its statistical significance (cf. Bouwhuis, Vedder, & Becker, 2015), to reflect the fact that the earlier an individual was born, the higher its lifespan could be, biasing our dataset. From these base models, we constructed a series of competing models which included SSTA in the natal year (the early-life-condition variable), whether it was the individual’s terminal reproductive event (1-0 variable), and the linear and quadratic expressions of age of first reproduction along with two-way interactions: SSTA in the natal year x age^2^, age of first reproduction x age^2^ and SSTA in the natal year x age of last reproduction. SSTA in the reproductive year was added to competing models for the analysis of offspring viability. Age of first reproduction was included to evaluate the indirect effects of early adversity, as this life history trait has been shown to vary with natal ENSO conditions (Ancona & Drummond, 2013) and is a major determinant of fitness (Pärt, 1995); its quadratic expression was added to detect an optimal onset of reproduction (e.g. Mourocq et al., 2016). The interactions were included to detect differences in senescence patterns due to natal conditions or age of first reproduction.

To test whether adult lifespan and lifetime reproductive success were influenced by early-life conditions, we used linear models (LMs). For both analyses, year of birth was also included in all competing models as a fixed variable. Variables included in the models were SSTA in the natal year and the linear and quadratic expression of age of first reproduction. For lifetime reproductive success adult lifespan was added to the competing models, along with its interaction with age of first reproduction.

To test for increased representation in warm water years of breeders of above average quality or peak reproductive ages (10-14 years, middle aged), or from cool water natal years (< −0.5 SSTA), as well as diminished representation of both young (1-9 years) and old (15-23 years) breeders, we built linear models including sex as a factor and using the linear expression of year of reproduction to account for progressive increase in the proportions of middle-aged and old birds among ringed birds with additional years of monitoring and ringing.

We compared the proportions of nestlings that fledge under different ENSO conditions by building a LM for each brood size. To compare the probabilities of recruiting after fledging in different ENSO conditions, and test for self-selection of above average breeders for reproduction in warm water conditions (self-selection), we built LMMs. For all three analyses, year of birth and SSTA in the natal year were included as fixed variables. For the recruitment analysis, body condition at fledging and chick rank (1^st^, 2^nd^ or 3^rd^ hatched) were included in the competing models, along with three two-way interactions: SSTA in the natal year x body condition, SSTA in the natal year x chick rank, and chick rank x body condition. Nest identity was added as a random effect because, in some cases, siblings were included in the analysis. Body condition and rank were included to test whether low values affect recruitment. The interactions tested whether poor natal environments increase the cost of poor body condition or low rank, and evaluated whether body condition at fledging varies with rank.

On the basis of AICc, ΔAICc, and AICc weights (*wi*) of the competing models, we selected the best-supported model, or averaged a subset of the best-supported models with a cumulative Akaike weight of ∼0.95. For each resulting model, we calculated the estimates (all reported effect sizes were standardized to two SDs) and 95% confidence intervals (CI). Model selection tables and summaries of best-supported and averaged models are provided in the supplementary materials.

## Results

### Fitness effects of natal ENSO conditions

Annual breeding success (number of fledglings produced) of 1893 female recruits from the 18 birth cohorts was unaffected by water temperature in the natal year (Table S2; Figure 1a). Natal water temperature affected the annual breeding success of 2143 male recruits, but not in the expected direction. Males that experienced warm natal waters produced slightly more fledglings annually than those born under more favourable conditions (β = 0.009 [95% Confidence Interval: 0.008 to 0.048]; Figure 1), with males fledged under a +0.8 °C SSTA (sea surface temperature anomaly) predicted to produce roughly 6% more fledglings than males fledged under a −0.8 °C anomaly (the extreme anomalies observed). Consistent with previous reports (Ancona et al., 2011; Drummond et al., 2003), boobies of both sexes increased their annual breeding success with earliness of laying and current coolness of the sea, and showed senescence after a peak in reproductive success at roughly ages 10-14 years (Beamonte-Barrientos, Velando, Drummond, & Torres, 2010; Alberto Velando, Drummond, & Torres, 2006). Importantly, natal SSTA did not interact with age (Table S3), showing that warm natal waters had no effect on either the rate of increase in breeding success during early adult years or the rate of decline during late adult years (reproductive senescence).

**Fig. 1.**
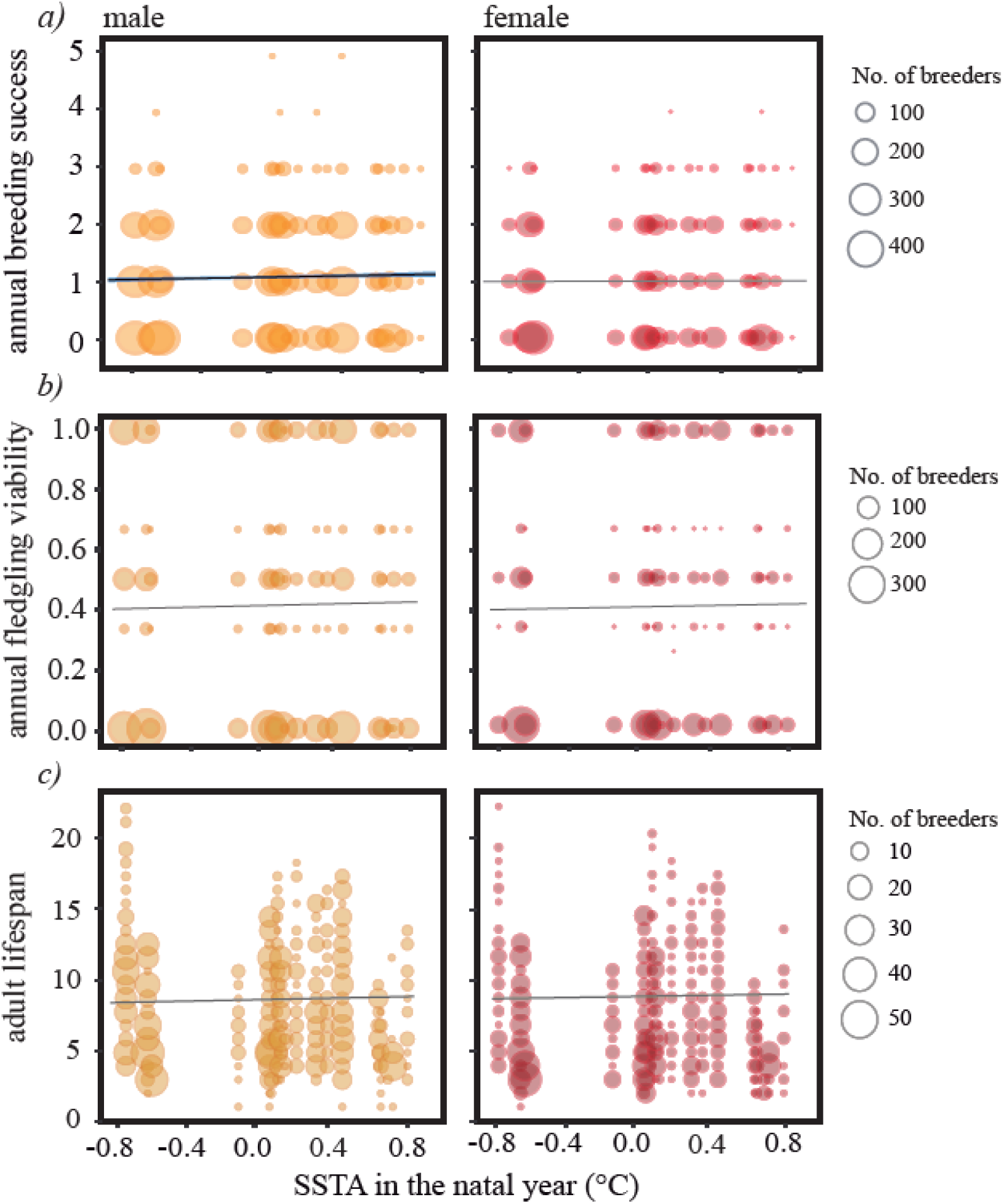
Effect of SSTA in natal year on annual breeding success, fledgling viability and lifespan. **a)** A male booby’s breeding success (fledglings produced) increased slightly with its natal SSTA (Sea Surface Temperature Anomaly), whereas a female’s breeding success was unaffected. Points represent 2143 male and 1893 female breeders observed over their lifespans. **b)** For 1646 male and 1329 female breeders, the proportion of their annually produced fledglings that recruited was unaffected by their natal SSTA. **c)** Adult lifespan of 1319 male and 1105 female breeders was not affected by natal SSTA. Solid line represents model predictions, with 95% confidence intervals in blue; grey lines show the statistically unsupported tendencies for the other five analyses.

Viability (probability of recruitment) of the fledglings produced by 1646 male and 1329 female recruits at all ages over the lifespan was unaffected by those recruits’ natal water temperature (Table S5; Figure 1b), although other variables were influential. Fledglings produced by both sexes in warm water years were less likely to recruit (Males: β = −0.194 [95% CI: −0.222 to −0.166]; Females: β = −0.195 [95% CI: −0.231 to −0.159]) and, at all ages, earliness of laying increased the viability of fledglings produced (Males: β = −0.128 [95% CI: −0.156 to −0.101]; Females: β = −0.112 [95% CI: −0.146 to −0.077]). A recruit’s age had a quadratic effect on the probability its offspring would recruit (Table S6), with peaks at roughly 10 years in males and 11 years in females.

The lifespan of recruits was unaffected by natal water temperature (Table S8; Figure 1c), but increased with late recruitment in both sexes (Males: 0.139 [95% CI: 0.090 to 0.0.189]; Females: 0.213 [95% CI: 0.160 to 0.266]), although this effect might arise simply because late recruits cannot die early.

Finally, lifetime reproductive success (total recruits produced) of both sexes was unaffected by natal water temperature (Table S11; Figure 2), although in both sexes age of recruitment and adult lifespan were influential. Boobies with younger ages of first reproduction produced more recruits over their lifetime than those that started later in life (Males: β = - 0.086 [95% CI: −0.126 to −0.046]; Females: β = −0.062 [95% CI: −0.119 to −0.013]). Greater adult lifespan, itself associated with delayed age of first reproduction, also increased lifetime production of recruits in both sexes (Males: β = 0.715 [95% CI: 0.670 to 0.761]; Females: β = 0.619 [95% CI: 0.561 to 0.677]). Note that despite achieving roughly 6% higher annual breeding success than cool natal water males over the lifespan, warm natal water males showed no corresponding increase in lifetime reproductive success, probably because they skipped more breeding seasons than cool water males (Ancona & Drummond, 2013). Warm natal water females showed no such increase in annual breeding success, but females are nearly one third heavier than males and more susceptible to food deprivation (Torres & Drummond, 1997; Velando, 2002).

**Fig. 2.**
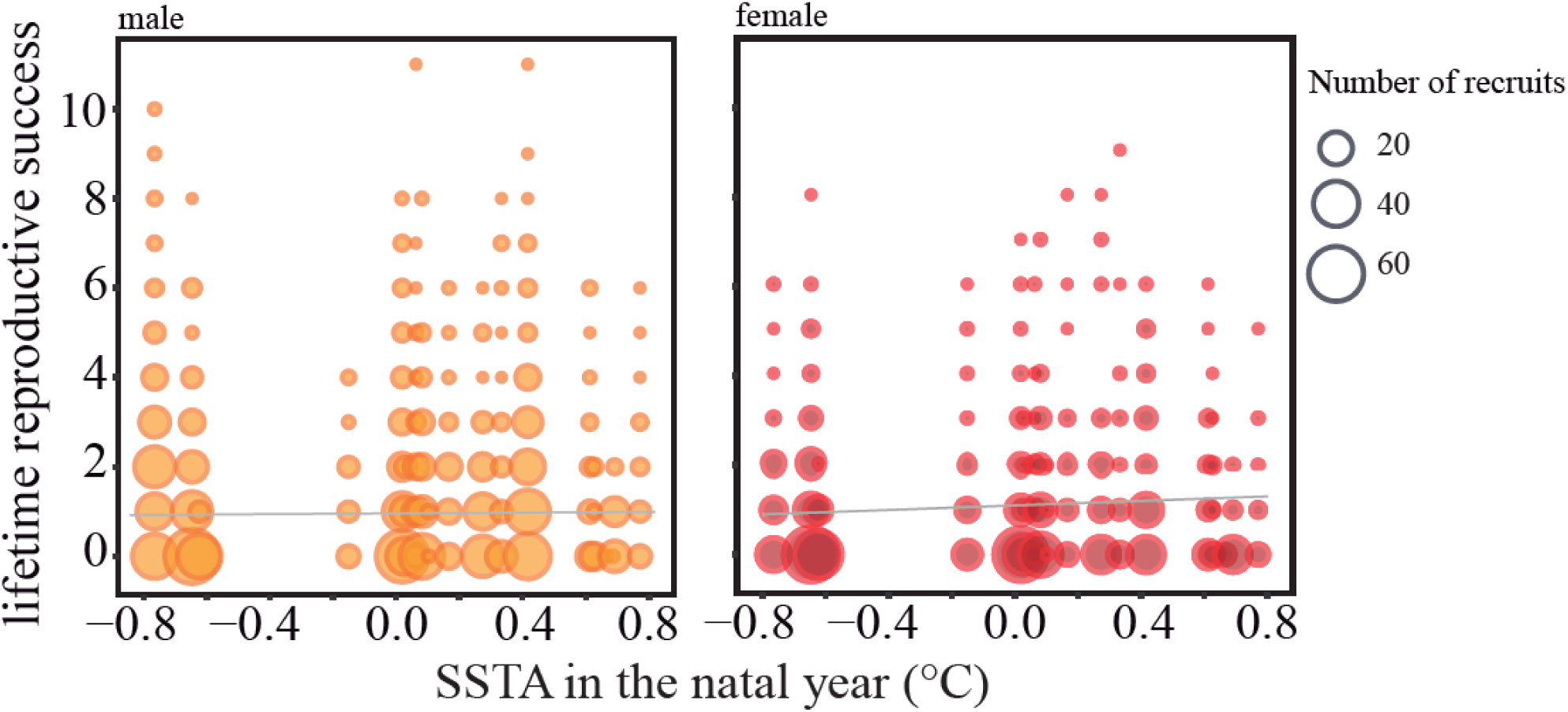
Lifetime production of recruits was unaffected by natal water temperature. Data for 1319 male and 1105 female breeders. Grey lines show the statistically unsupported tendency for production to increase in both sexes with warm natal water.

### Self-selection for breeding in El Niño conditions

In warm water years neither boobies of above average quality nor boobies born in cool water (La Niña) years increased their proportional representation among breeders (n = 4036; Tables S13-S15) but, contrary to expectation, middle aged boobies substantially diminished theirs (β = −0.317 [95% CI: −0.470 to −0.164]) and the proportions of both young (1-9 years) and old (>14 years) boobies among breeders increased (Young: β = 0.193 [95% CI: 0.068 to 0.318]; Old: β = 0.175 [95% CI: 0.020 to 0.331]).

In this booby, despite young and old individuals showing inferior breeding success (see fitness effects), young-with-old parental pairings produce the most viable (likely to recruit) fledglings (Drummond & Rodríguez, 2015), so we hypothesized that overrepresentation of young and old breeders in warm years could increase the proportion of young + old pairings and hence the average quality of fledglings and recruits. We tested for this effect on pairings by calculating the proportional representation of all six possible pairings of young, middle aged and old boobies in every year between 1993 and 2017 (Table S16). Sex, as a factor, was not included in the LMs because greater viability of offspring of young-old pairs is independent of which sex is the older pair mate (Boyko, 2008; Drummond & Rodríguez, 2015). These analyses showed that warm water is indeed associated with an increase in the proportion of young + old pairings, and also with a decrease in the proportion of young + middle aged pairings (Young + old: β = 0.280 [95% CI: 0.072 to 0.487]; Young + Middle aged: β = −0.340 [95% CI: −0.614 to −0.067]; Figure 3).

**Fig. 3.**
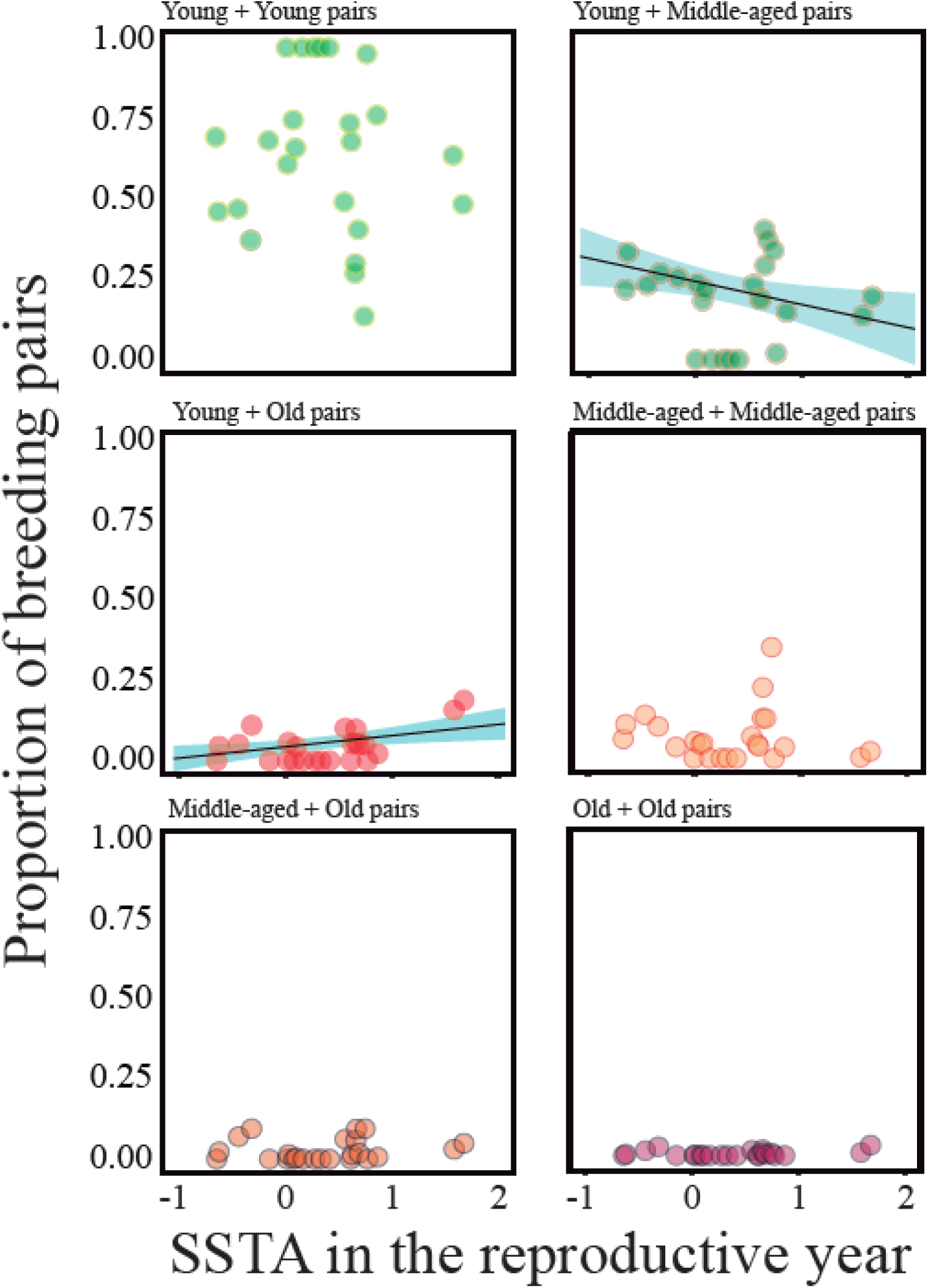
In warm water years the proportion of young + old pairs increased, and the proportion of young + middle aged pairs decreased.

### Viability selection in El Niño conditions

Both nestling and juvenile mortality increased with the warmth of natal waters, potentially increasing the influence of any recruitment bias in favour of higher quality cohort members. Nestlings from all three brood sizes were less likely to fledge in the warmest natal condition than in the coolest (Table S17;Fig 4a), with a greater differential in three-chick broods (β = −0.27 [95% CI: −0.32 to −0.22]) than in one- and two-chick broods (β = −0.10 [95% CI: −0.12 to −0.08] and β = −0.14 [95% CI: −0.16 to −0.12], respectively), and with 2^nd^ and 3^rd^ hatched chicks especially liable to fail, independent of natal water temperature (Table S19; Figure. 4b). Fledgling body condition was key: in all cohorts, poor condition reduced the probability of recruiting (β = 0.07 [95% CI: 0.05 to 0.08]; Figure. 4c). Importantly, compared to cohorts from the coolest water, 23% fewer chicks from cohorts of the warmest water fledged, and 26% fewer fledglings from cohorts of the warmest water went on to recruit.

**Fig. 4.**
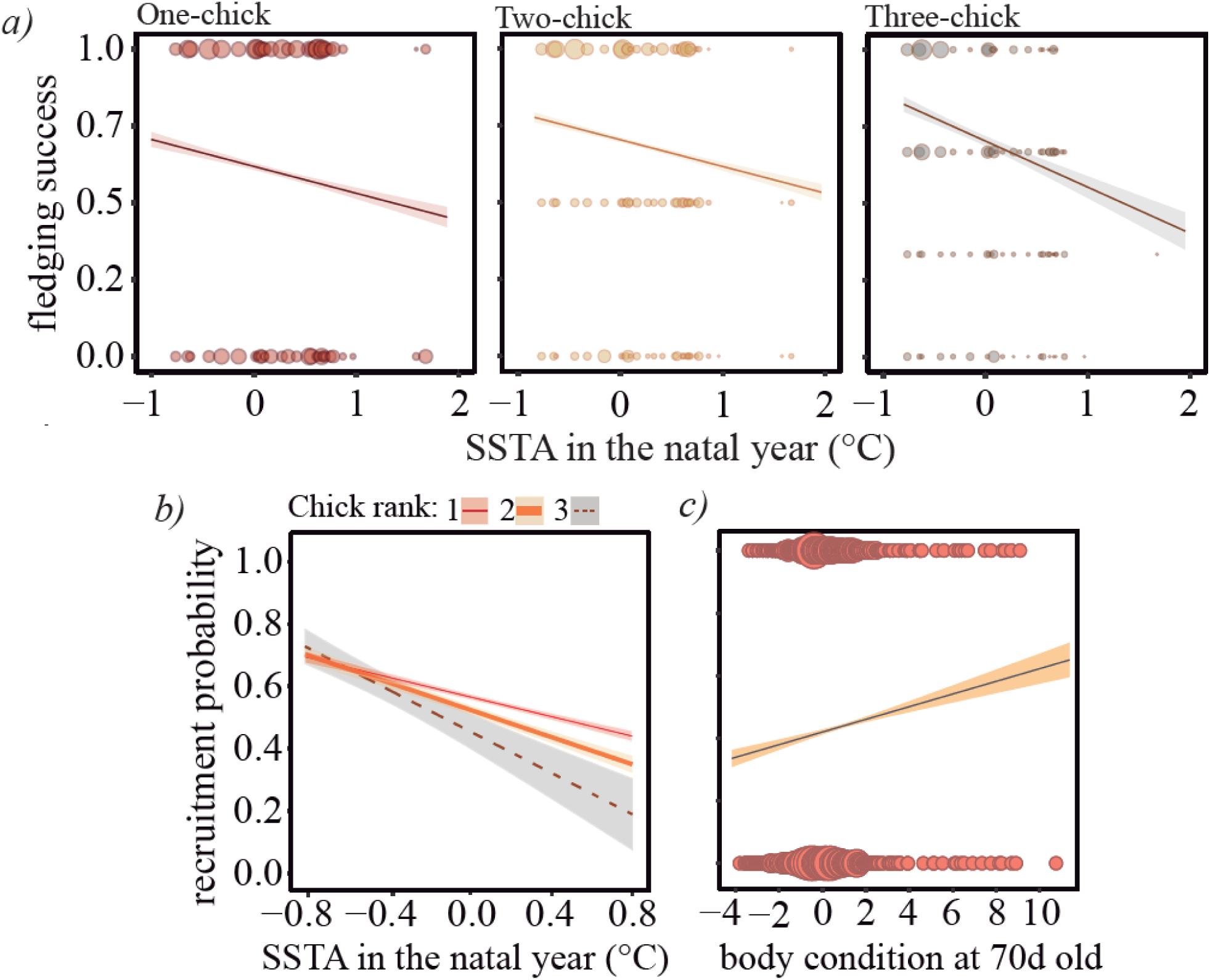
a) The proportion of chicks from 15621 broods that managed to fledge decreased with the warmth of natal waters. The probability of recruiting for 10604 unsexed fledglings b) decreased with the warmth of natal waters, especially for second and third chicks, and c) in every year was higher for offspring that fledged with good body condition.

## Discussion

Given the developmental susceptibility of birds to experimental stresses in infancy, including food scarcity, compensatory growth, elevated corticosterone and parasitic infection, as well as poor weather, habitats, prey availability, parental care and late fledging (Drummond & Ancona, 2015), it is remarkable that the annual breeding success, offspring viability, adult lifespan and lifetime reproductive success of adult booby generations are not affected by exposure to El Niño in their natal year. However, it remains to be shown that the viable offspring of recruits from warm water cohorts pay no reproductive costs after recruitment (an intergenerational effect; Naguib & Gil, 2005; cf. Bouwhuis et al., 2015). We conclude that, even though warm waters can severely prejudice breeding participation of the focal population and impact not only clutch sizes but also hatching, fledging and recruitment success (Ancona, Zúñiga-Vega, Rodríguez, & Drummond, 2018), as well as adult survival (Oro et al., 2010), the recruits derived from warm water cohorts can contribute normally to the population’s growth rate.

We identified two mechanisms that probably contribute to warm water recruits performing as well as cool water recruits. Self-selection for breeding in warm water years probably makes a small contribution. High-quality boobies, boobies raised in cool water conditions, and boobies of prime reproductive ages (middle aged) do not increase their representation among breeders in warm years. However, young + old pairs increase their representation to 7% of breeding pairs in El Niño years, so their progeny could boost the average quality of recruits from El Niño years provided the high quality conferred on fledglings by those pairings persists into adult life. Considering the extremes, a fledgling produced by a pair 1yr + 21yr old is roughly 80% more likely to recruit than that of a pair 1yr + 1yr old (Drummond & Rodríguez, 2015). Nonetheless if, as hinted in Figure 3, the relationship between water temperature and proportion of young + old pairings is a step function rather than a linear function, then that representation may ascend to roughly 20% of breeding pairs in El Niño years, implying a greater potential boost to average recruit quality. Note that increases in the proportions of young and old breeders in El Niño years could serve functions unrelated to the benefit of pairing with opposite-age partners. For example, red-footed booby (*S. sula*) pre-breeders show preference for recruiting in El Niño years and are thought to do so because competition may be weaker (Cubaynes et al., 2010), and after age 14 years blue-footed boobies may be penalized for skipping reproduction because their annual survival is in decline and increasingly variable (Drummond, Rodriguez, & Oro, 2011).

The second mechanism, viability selection on offspring, is more prevalent and probably more influential than filtering of breeders. The increase in mortality among booby offspring in warm water years should raise the average reproductive value of recruits to the extent that mortality falls disproportionately on low quality individuals. In the nestling period, dominance ranks in the brood derive from 4-day hatch intervals and selective mortality in broods of two and three is dictated by hatch order rather than individual quality (Drummond & Garcia Chavelas, 1989; Drummond, González, & Osorno, 1986), so viability selection may be weak. By contrast, in the pre-breeding period, when juvenile boobies transition to independent feeding and compete for reproductive opportunity in the colony, their survival and recruitment, associated with high body condition at fledging, may depend largely on individual quality. If so, the increase in fledgling mortality from roughly 25% in cohorts from the coolest years to 60% in cohorts from the warmest years marks an increase in viability selection that could substantially raise the average phenotypic quality of recruits from El Niño cohorts.

The explanation for reproductive equivalence of generations of recruits from El Niño cohorts lies in a combination of developmental resilience of blue-footed booby offspring, increased viability selection among them, especially during the pre-breeding period, and increased participation of young + old breeding pairs. Developmental resilience probably depends on increased investment by parents caring for late, slow-growing offspring during food scarcity, and also on physiological adaptations and life history adjustments of those offspring, which tend to recruit at an early age and skip more breeding years (Ancona & Drummond, 2013). We found that early recruitment generally increases lifetime reproductive success, and warm water cohorts obtain this benefit while avoiding both increase in mortality at any adult age (Ancona et al., 2018) and the longevity cost (Table S12) of reproductive investment early in life, apparently by skipping some breeding seasons. Blue-footed boobies may not be alone in this ability to parry impacts of a stressful natal environment by making life history adjustments. Females of the reintroduced Mauritius kestrel (*Falco punctatus*) mitigate fitness penalties of an anthropogenically altered natal environment by modifying their schedule of reproductive effort (Cartwright et al., 2014), although it is unclear whether their lifespan is affected.

The prevalence among animal species of El Niño cohort effects on the reproductive value of recruits is unknown, and we may find that many species have evolved ways of coping. Exposure of species of the southern and eastern Pacific ocean to ENSO-like oscillations since at least the mid-Holocene (Corrège et al., 2000) has probably honed the evolution of developmental, behavioural and life history traits that contribute to the reproductive equivalence of adult generations. Nonetheless, given the well-documented developmental susceptibility of infant vertebrates to adverse natal environments and the increasing severity of El Niño events, it is likely that natal El Niño conditions compromise the reproductive potential of affected generations of some species, and that such effects will increase in the near future. We should examine other species, to document natal El Niño impacts on reproductive potential, to identify filtering effects and other means by which species sidestep or resist impacts, and to explore their demographic consequences.

## Acknowledgments

We thank numerous volunteers for enthusiastic work in the field and on the database; the Armada de México, staff of SEMARNAT and local fishermen for logistical support. Santiago Ortega was supported by grants from the Consejo Nacional de Ciencia y Tecnología (CONACYT) and the Coordinación General de Estudios de Posgrado, UNAM, during his studies in the Posgrado de Ciencias Biológicas, UNAM. Funding for this project was provided by grants to HD from CONACYT (255826) and the National Geographic Society (9974-16). For comments on earlier drafts we thank María Valverde, Alejandro Gonzalez-Voyer and Sergio Ancona. This paper constitutes the fulfilment of a requirement for obtaining the degree of Maestro en Ciencias Biológicas in the field of Biología Evolutiva of the Posgrado en Ciencias Biológicas de la UNAM.

## Author’s contributions

SO and HD conceived the study, HD and CR collected the data, SO analysed the data, SO and HD led the writing of the manuscript. All authors contributed critically to the drafts and gave final approval for publication.

## Data accessibility

Data files are available from Dryad upon manuscript acceptance.

## Supporting information

Addition Supporting Information may be found in the online version of this article.

